# *Rickettsia rickettsii* encodes a secretory lipase that facilitates intracytosolic colonization in host cells

**DOI:** 10.1101/2024.09.16.613323

**Authors:** Mohammad Sadik, Imran Moin, Saif Ullah, M. Sayeedur Rahman, Oliver H. Voss

**Author notes:** Correspondence (O.H.V.).

## Abstract

The key cellular processes required for rickettsial obligate intracellular lifestyle, include internalization by phagocytosis, regulation of intracellular trafficking, and evasion of lysosomal destruction to establish an intracytosolic replication niche, remain poorly defined. Recent reports showed that rickettsial phospholipases play an important role in vacuolar escape, but their functions are dispensable depending on the host cell-type. Here, we report the identification of a highly conserved putative lipase containing a Serine hydrolase motif (GXSXG), named RLip (*<u>R</u>ickettsia* <u>Lip</u>ase). Our work reveals that RLip expression is cytotoxic to yeast cells, a genetically tractable heterologous model system. We demonstrate that RLip possesses lipase enzymatic activity and show a lipid specificity towards phosphoinositide (PI)(3), PI(3,4,5)P_3_, and PI(3,4)P_2_, and to a lesser extent PI(4,5)P_2_. Further, we found that RLip expression is induced during infection of pathogenic *R. rickettsii*, while its expression is low or undetectable for *R. parkeri* (mild-pathogenic) and *R. montanensis* (non-pathogenic), respectively, during host invasion. Intriguingly, RLip is highly enriched in the cytoplasmic fraction of host cells, however, minimally retained by the rickettsiae themselves, suggesting RLip is synthesized during infection and then secreted into the host cell cytoplasm. Neutralization of RLip activity, by antibody-blocking, significantly abrogated *R. rickettsii* escape from bactericidal phagolysosomal fusion, suggesting RLip plays a critical role in facilitating the intracytosolic colonization of pathogenic *R. rickettsii*.

**Importance:** Arthropod-borne rickettsial diseases are on the rise globally, presenting a perilous threat to humans and livestock. However, our inadequate understanding on how *Rickettsia* manipulates cellular processes, including the evasion of lysosomal destruction, has impaired the development of effective therapeutic interventions.

Here, we identify of a conserved putative lipase containing a Serine hydrolase motif, named RLip (*<u>R</u>ickettsia* <u>Lip</u>ase). Our work demonstrates that RLip possesses lipase enzymatic activity and is enriched in the cytoplasm of host cells, while minimally retained by the bacteria itself. Neutralization of RLip activity abrogated *R. rickettsii* escape from bactericidal phagolysosomal fusion. In sum, our data support a mechanism by which pathogenic *R. rickettsii* employs RLip to escape from bactericidal phagolysosomal fusion in order to colonize the host.

## Introduction

Rickettsioses are vector-borne diseases, presenting a perilous threat to humans and livestock. In fact, tick- and flea-borne rickettsial diseases are on the rise globally and our inadequate understanding on how *Rickettsia* interacts with their mammalian host has impaired the development of effective interventions against pathogenic rickettsial infections (1).

Rickettsiae infection begins at the bite site of the host skin, transmitted by their specific arthropod vector. During invasion, rickettsiae can manipulate various cellular processes, including host membrane dynamics, actin cytoskeleton, phosphoinositide (PI) metabolism, intracellular signaling and immune defense responses. In fact, as strict obligate intracellular Gram-negative bacteria, rickettsiae are required to escape from the destruction of phagolysosomal fusion into a metabolite-rich host cytosol to establish a replication niche and ultimately disseminate into neighboring cells/organs of the host(2–6). To orchestrate such a complex cellular task, *Rickettsia* and other intracellular bacteria, utilize an arsenal of effectors, including membranolytic enzymes. For instance, *Listeria monocytogenes* uses the cholesterol-dependent cytolysin listeriolysin O (LLO) and several phospholipase C enzymes to promote vacuolar escape(7–11), while *Pseudomonas aeruginosa* releases ExoU, a phospholipase A_2_ (PLA_2_), to accomplish their pathogenic life cycle(12, 13). In addition, *Legionella pneumophila* secrets two phospholipases, VipD and VpdC, to facilitate their intracellular lifestyle(14, 15), while *Shigella flexneri* utilizes the IpaB and IpaC invasins to promote membrane rupture and host invasion(16–18). In the case of rickettsiae, we reported that rickettsial phospholipase A_2_ enzymes Pat1 (present in all rickettsiae), and Pat2 (variably present) were involved in the phagosomal escape to support host colonization(19–21). Also, reports from other laboratories revealed that phospholipase D (PLD) (present in all rickettsiae)(22) and Pat1(23) play a role in vacuolar escape, however, theses enzymes have also been reported to be dispensable depending on the host cell type(22, 23). Given these reports and the fact that *Rickettsia*, as an obligate intracellular parasite, requires membranolytic enzymes to escape from host lysosomal destruction, we hypothesize that additional rickettsial effectors regulate host membrane dynamics to facilitate vacuolar escape into the host cytosol. In this effort, our bioinformatical analysis on the rickettsial genomes(24) resulted in an identification of a putative lipase, named RLip (*<u>R</u>ickettsia* <u>Lip</u>ase), that harbored a highly conserved Serine hydrolase motif (GXSXG). Functional characterization revealed that recombinant RLip of *R. rickettsii* possesses lipase enzymatic activity, and its expression is cytotoxic to yeast cells. Anti-RLip antibody-mediated neutralization of RLip only moderately affected *R. rickettsii* infection of host cells, however, it abrogated significantly rickettsial escape from lysosomal fusion suggesting that RLip plays a critical role in facilitating the intracellular survival of pathogenic *R. rickettsii*.

## Results

### RLip is a secreted effector with a putative lipase domain

As an obligate intracellular parasite, *Rickettsia* requires membranolytic enzymes to escape membrane-bound vacuoles, to gain access to the metabolites rich host cytosol. Recent reports from others and our laboratory have demonstrated the role of rickettsial membranolytic effectors in facilitating colonization in host cells(19, 20, 22, 23, 25, 26). However, the redundancy and sporadic presence of these rickettsial effectors(21–23) suggest the presence of additional membranolytic enzymes required for disrupting the vacuolar membrane to mediate the escape from lysosomal destruction during the rickettsial invasion process. To test this hypothesis, we searched the rickettsial genomes using the web-based Phyre2 program(24) and identified a putative secretory lipase with a Serine hydrolase motif (GXSXG), named RLip. Sequence alignment of RLip with phospholipases, including ExoU (*Pseudomonas aeruginosa*)(12); VipD (*Legionella pneumophila*)(14), VpdC (*Legionella pneumophila*)(15), and rickettsial Pat1 as well as Pat2(19, 20), depicted a highly conserved Serine hydrolase motif (**Fig. 1A**). Further comparison shows that the Serine hydrolase motif was highly conserved amongst RLip molecules from other *Rickettsia* species (up to ∼95% identity) (**Fig. 1B**). As lipases belong to the α/β hydrolase family with an active site Serine situated at the catalytic loop between α-helices and a β-sheet(27–29), we generated a homology model of RLip using the Phyre2 software(30). Our model depicted that RLip harbors a classical lipase structure consisting of a β-sheet surrounded by two α-helices (Fig. S1A) with an active site Serine placed in the catalytic loop (Fig. S1A; inset). Additional comparison of the lipase structure with other phospholipases (ExoU, VipD, VpdC, Pat1, as well as Pat2) provided further evidence that RLip harbors a highly conserved lipase motif (Fig. S1B). Of note, all presented models were validated by Ramachandran plot using the RAMPAGE software(31).

**Fig. 1.**
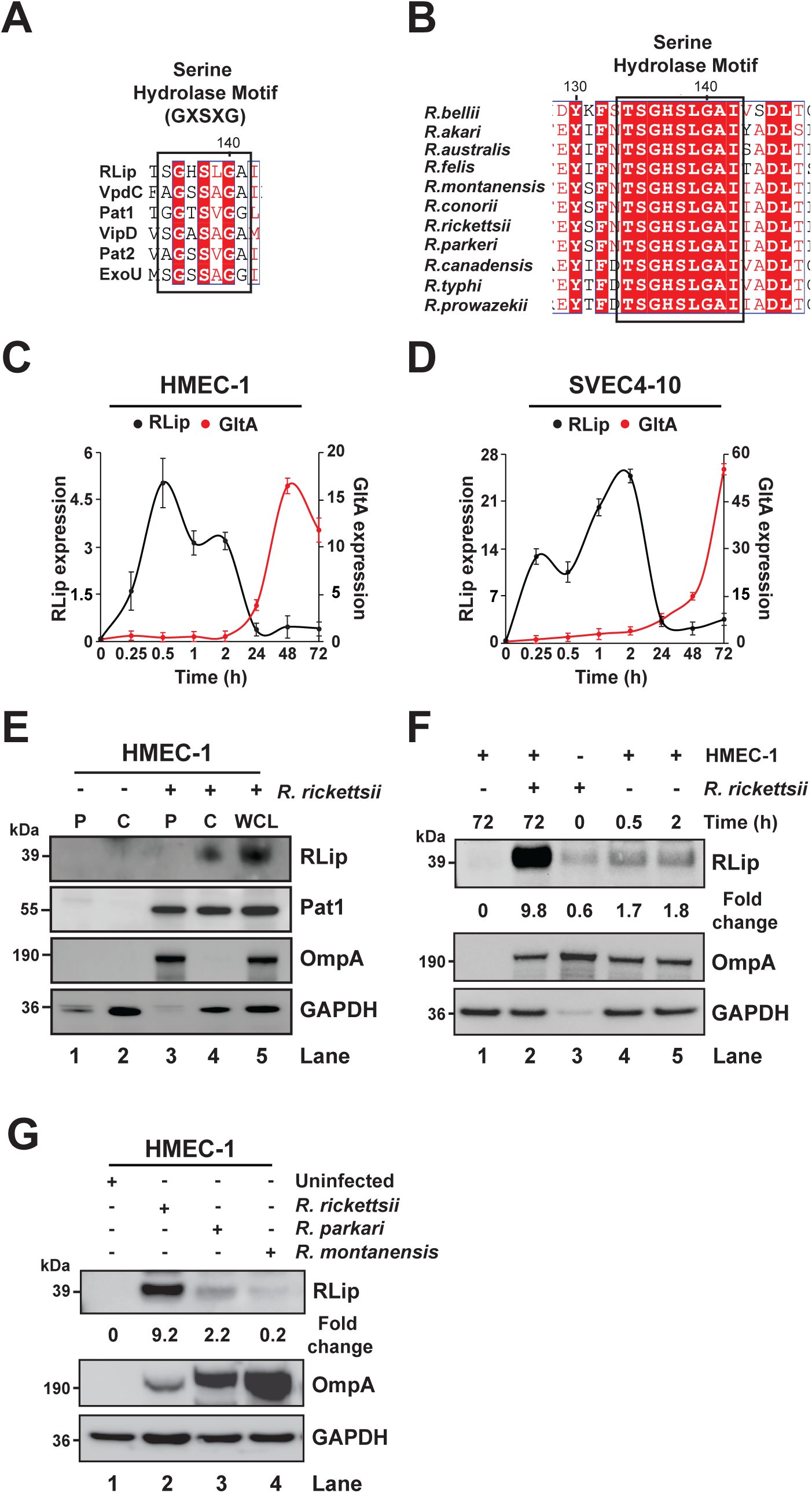
RLip is a secreted effector harboring a conserved hydrolase motif. **(A)** Comparative sequence alignment of catalytic Serine hydrolase motifs among rickettsial proteins [Pat1, Pat2, and *Rickettsia* Lipase (RLip, locus_tag # A1G-01170)], and other bacterial lipases VipD (*Legionella pneumophila*), VpdC (*Legionella pneumophila*), and ExoU (*Pseudomonas aeruginosa*). Highly conserved Serine hydrolase motif amino acids (GXSXG) are highlighted in red. (**B**) Sequence alignment of the conserved Serine hydrolase motif (GXSXG) of *R. rickettsii* RLip with other *Rickettsia* species. (**C-D**) Expression kinetics of RLip (black line) and rickettsial citrate synthase (*GltA,* red line) during *R. rickettsii-*infection of HMEC-1 (**C**) and SVEC 4-10 (**D**) was determined by RT-qPCR. Rickettsial growth was determined by *GltA* expression. Both *RLip* and *GltA* expression was normalized with respect to *GAPDH* expression as described previously(51, 57). (**E**) Uninfected or *R. rickettsii*-infected HMEC-1 cells were lysed with 0.1 % Triton X-100 and separated into cytoplasmic (C) and pellet (P) fractions. Samples were immunoblotted with anti-RLip, anti-Pat1 (positive secreted *R. rickettsii* effector control), anti-OmpA/B (as control for *R. rickettsii* surface protein), or anti-GAPDH Abs (host cytoplasmic control protein). Whole cell lysates (WCL) were used as expression control for all target proteins. (**F**) Partially purified rickettsiae were incubated with HMEC-1 cells for 0.5 or 2 hrs at 34°C and analyzed by immunoblotting using anti-RLip, anti-GAPDH, and anti-OmpA/B Abs. Uninfected and *R. rickettsii-*infected WCL (72 hrs) were used as expression controls. (**G**) HMEC-1 cells were infected with spotted fever group (SFG) rickettsiae, including *R. rickettsii* (highly pathogenic), *R. parkeri* (moderate pathogenic) and *R. montanensis* (non-pathogenic) for 72 hrs. Samples were analyzed by western blot analysis using anti-RLip, anti-GAPDH, and anti-OmpA/B Abs. Densitometry in panels **F**, and **G** was performed using Fiji software, and data are presented as fold change between the ratios of RLip/OmpA. Error bars (**C**, **D**) represent means ± standard error of the mean (SEM) from three independent experiments. Images in panels **E**, **F**, and **G** are a representative of 3 independent experiments.

Given that phospholipases have been shown to play an important role during host cell infection(19, 20, 22, 23, 25, 26), we first evaluated the transcriptional pattern of RLip during *R. rickettsii-*infection of various mammalian cells (HMEC-1, Vero76, and SVEC 4-10) by RT-qPCR. Our data demonstrated that transcription levels of RLip expression were increased as early as 15 min post-infection and remained elevated up to 2 hrs [**Figs. 1C**, and D, and Fig. S2A), while levels of rickettsial growth (as depicted by normalized *GltA* expression) appeared to increase at 24 hrs post-infection [**Figs. 1C**, and D, and Fig. S2A). These findings indicate that RLip likely plays a critical role during the host invasion. To define the function of RLip during host infection, we raised an antibody against RLip protein (anti-RLip) and determined its specificity using *R. rickettsii*-infected Vero cells and recombinant RLip protein (Fig. S2B). Next, we examined whether RLip is secreted from *R. rickettsii* during infection of HMEC-1 cells by performing cellular fractionation into cytoplasmic(C) and pellet(P) fractions. The cytoplasmic [(C), carrying host soluble and rickettsial secreted proteins] and pellet [(P), carrying rickettsiae and insoluble host debris] fractions were analyzed by immunoblotting as described previously(32). We observed that glyceraldehyde-3-phosphate dehydrogenase (GAPDH; host cytoplasmic protein) appeared in the cytoplasm of both uninfected and infected cells (**Fig. 1E**, lanes 2 and 4). Of note, the observed faint GAPDH bands within both pellet fractions are likely the result of incomplete lysis of the host cells or residual supernatants left with the pellet fractions (**Fig. 1E**, lanes 1 and 3). OmpA, a rickettsial outer membrane protein(32, 33), was only present in the pellet fraction of infected cells (**Fig. 1E**, lane 3), suggesting that lysis of host cells in the presence of 0.1% Triton X-100 is not affecting the cell surface integrity of *R. rickettsii*. Furthermore, Pat1 was present in both the pellet (**Fig. 1E**, lane 3) and cytoplasm (**Fig. 1E**, lane 4) of infected cells, implying that Pat1 is partially secreted into the host cell cytoplasm during infection. For RLip, the protein was secreted into the cytoplasm of infected HMEC-1 cells and was not retained to a significant level by the bacteria itself (**Fig. 1E**, lane 4 vs. 3). Of note, similar findings of the secretion pattern for RLip and Pat1 were observed during *R. rickettsii* infection of Vero76 cells (Fig. S2C). Due to its intriguing expression and secretion pattern, we tested the hypothesis whether RLip expression was induced upon host cell invasion. In this effort, we infected HMEC-1 cells for 0.5 and 2 hrs with partially purified *R. rickettsii* from host cells and analyzed RLip protein expression by western blot analysis. Our data revealed that RLip expression was rapidly induced, around 3-fold, after 0.5 and 2 h post infection (**Fig. 1F**, lane 3 vs. 4 or 5). Next, we compared the protein expression profile of RLip during host cell infection among spotted fever group (SFG) rickettsiae, including *R. rickettsii* (highly pathogenic), *R. parkeri* (moderate pathogenic) and *R. montanensis* (non-pathogenic), by western blot analysis. Our findings showed that RLip was highly expressed during *R. rickettsii* infection, while its expression was low or undetectable during *R. parkeri* and *R. montanensis* infection, respectively (**Fig. 1G**). In sum, these data suggest that RLip is expressed and efficiently secreted into the cytoplasm of the host cells and may function as a potential effector molecule required for invasion of highly pathogenic *Rickettsia* species.

### RLip possess lipase and cytotoxic activities

Our bio-informatic analysis predicted RLip as a putative lipase with a conserved Serine hydrolase motif (GXSXG) (**Fig. 1**). To further interrogate the functional characteristics, we generated codon optimized full-length wild-type His-tagged recombinant RLip protein (rRLip-WT-CO) as well as a catalytic active site mutant, rRLip-S138A-CO, in which a Serine (S) was mutated to an Alanine (A) at position 138 (**Fig. 2A**). As previously demonstrated for Pat1 and Pat2(19, 20, 23), our data reveal that rRLip-WT-CO possesses lipase enzymatic activity, while mutagenesis of the S138 residue significantly reduced the activity of RLip (**Fig. 2A**). Of note, heat-inactivation (Hi) of either rRLip-WT-CO or rRLip-S138A-CO resulted in the loss of lipase activity (**Fig. 2A**). To test if host cell factor(s) contribute to the lipase activity of RLip, we measured the enzymatic activities of rRLip-WT-CO or rRLip-S138A-CO in the presence of untreated lysates from human endothelial cells (HMEC-1) and found that the addition of lysate enhanced the lipase activity of rRLip-WT-CO but not that of rRLip-S138A-CO (**Fig. 2A**). These findings suggest that RLip possesses lipase activity, which is enhanced by host cell factor(s).

**Fig. 2.**
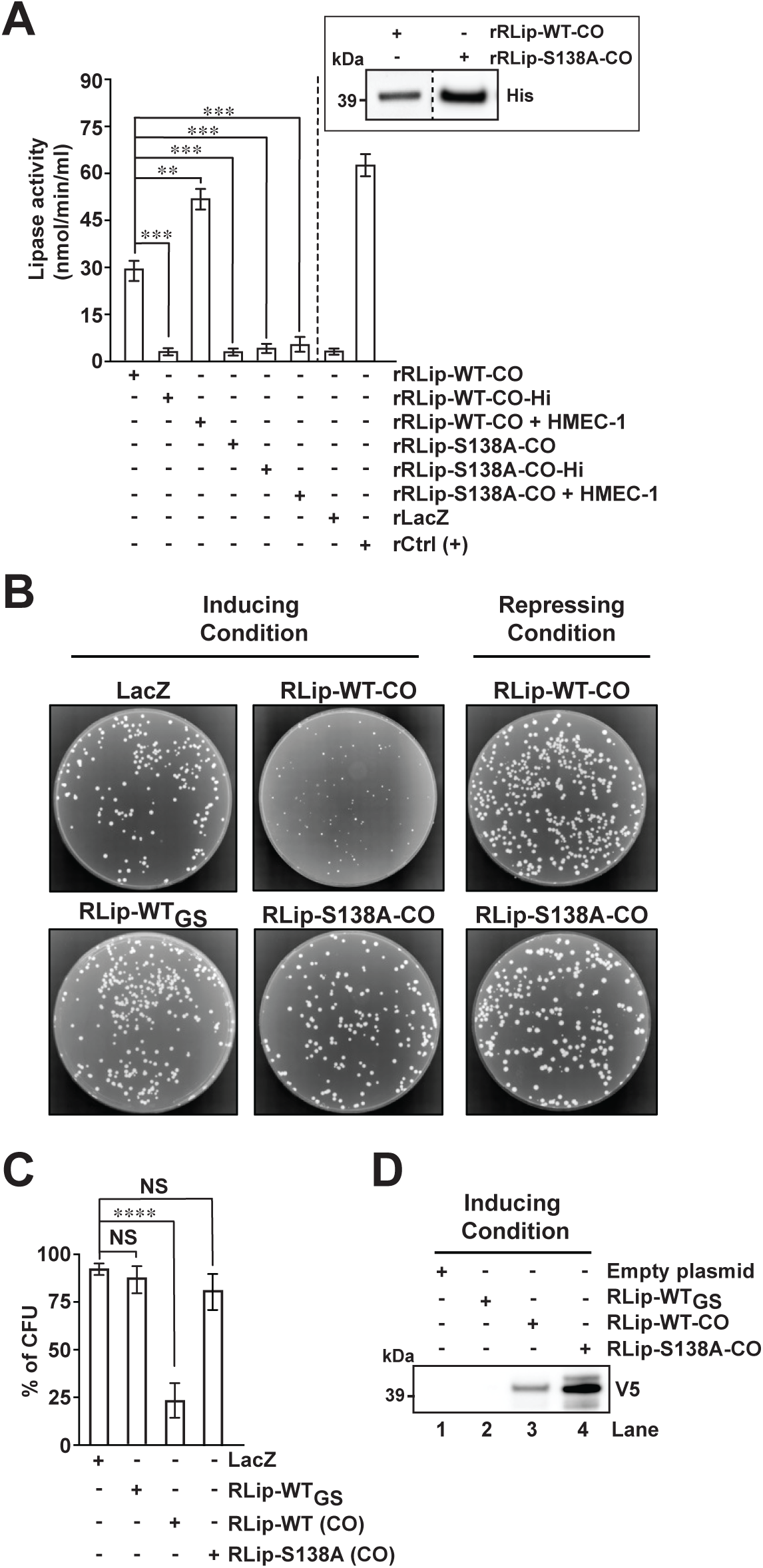
RLip is a rickettsial effector with lipase and cytotoxicity activities. (**A**) Lipase activity of purified codon-optimized recombinant(r) rRLip-WT-CO, rRLip-S138A-CO and heat-inactivated rRLip-WT-CO-Hi, rRLip-S138A-CO-Hi proteins in the absence or presence of host cell lysates (HMEC-1) was assessed as described previously(19, 20). The recombinant (r) LacZ protein and a lipase derived from *Chromobacterium* were used as a non-specific protein and positive (+) control, respectively. Inset represents a western blot analysis of the utilized rRLip-WT-CO and rRLip-S138A-CO using an anti-His Ab. (**B**) Transformed yeast cells were streaked onto inducing (SC-U+Gal) or repressing (SC-U+Glu) agar and incubated at 30°C for 3 days. (**C**) Cytotoxicity assay in yeast strain INVSc1 transformed with plasmids expression RLip-WT_GS_, RLip-WT-CO or lipase mutant RLip-S138A-CO was performed as described previously(19, 20). LacZ plasmid was used as control in panels **B**, and **C**. (**D**) Western blot analysis of RLip-WT_GS_, RLip-WT-CO or lipase mutant RLip-S138A-CO expression in yeast strain INVSc1 under inducing conditions (SC-U+Gal medium). The total proteins from yeast cells carrying the appropriate plasmid were probed with anti-V5 Ab. Error bars in panels **A**, and **C** represent means ± SEMs (standard errors of the means) from 3 independent experiments; NS, not significant; \*\**P* ≤ 0.01; \*\*\**P* ≤ 0.005; \*\*\*\**P* ≤ 0.001. Images in **A**, **B**, and **D** are a representative of 3 independent experiments.

It is well-established that heterologous model systems are an effective tool to assess biological functions of virulence factors. In fact, work from others and our laboratory have used *S. cerevisiae* as a genetically tractable system to demonstrate the cytotoxic effects of various effectors, including among others ExoU (*P. aeruginosa*)(12, 34), Pat1, and Pat2 (*Rickettsia*)(19, 20). To assess the cytotoxicity of RLip, we cloned codon optimized RLip-WT-CO and the catalytic active site mutant, RLip-S138A-CO, into the pYES2/CT vector and performed the yeast cytotoxicity assay as previously described(19, 20). Of note, pYES2/ RLip-WT_GS_ (carrying RLip encoded by *R. rickettsii* wild-type genome sequence (GS)] and pYES2/CT/LacZ plasmids were used as controls. Yeast transformants carrying either pYES2/CT/LacZ or pYES2/RLip-WT_GS_ grew well under either inducing (SC-U+Gal) or repressing conditions (SC-U+Glu) (**Fig. 2B**, and **C**). However, yeast carrying pYES2/RLip-WT-CO showed a significant reduction in growth under inducing condition (SC-U+Gal), while yeast carrying pYES2/RLip-S138A-CO showed no growth inhibition (**Fig. 2B**, and **C**). Further evaluation of RLip-WT-CO and RLip-S138A-CO protein expression within the yeast transformants by western blot analysis confirmed that both wild-type and mutant RLip proteins were expressed (**Fig. 2D**). Collectively, our data suggest that RLip harbors both lipase and cytotoxic activities, which requires a functional Serine hydrolase motif.

### RLip preferentially targets PI(3)P, PI(3,4,5)P_3_, and PI(3,4)P_2_ lipids

Membranolytic enzymes, like Pat1, and Pat2, target host lipids to promote escape from membrane-bound vacuoles to facilitate host cell invasion to establish an intracytosolic niche(19, 20, 23). Given these prior reports on Pat1, and Pat2, as well as our current data showing RLip possesses lipase enzymatic activity (**Fig. 2**), we examined if RLip targets phosphoinositides (PIs) using HeLa cells co-expressing green fluorescent protein (GFP)-tagged PI biosensors(32, 35, 36) with either pcDNA4-Flag empty vector, pcDNA4-Flag-RLip-WT, or pcDNA4-Flag-RLip-S138A. Strong colocalization was observed between RLip-WT and the biosensors for phosphatidylinositol 3-phosphate [PI(3)P], phosphatidylinositol 3,4-bisphosphate [PI(3,4)P_2_], and phosphatidylinositol 3,4,5-trisphosphate [PI(3,4,5)P_3_], whereas only a weaker correlation between RLip-WT and the phosphatidylinositol 4,5-bisphosphate [PI(4,5)P_2_] sensor was detected (**Fig. 3**). In contrast, overexpression of RLip-S138A resulted in a detectable but overall non-significant reduction in colocalization with all tested PI sensors (**Fig. 3**) suggesting that RLip but not its lipase activity is involved in substrate recognition.

**Fig. 3.**
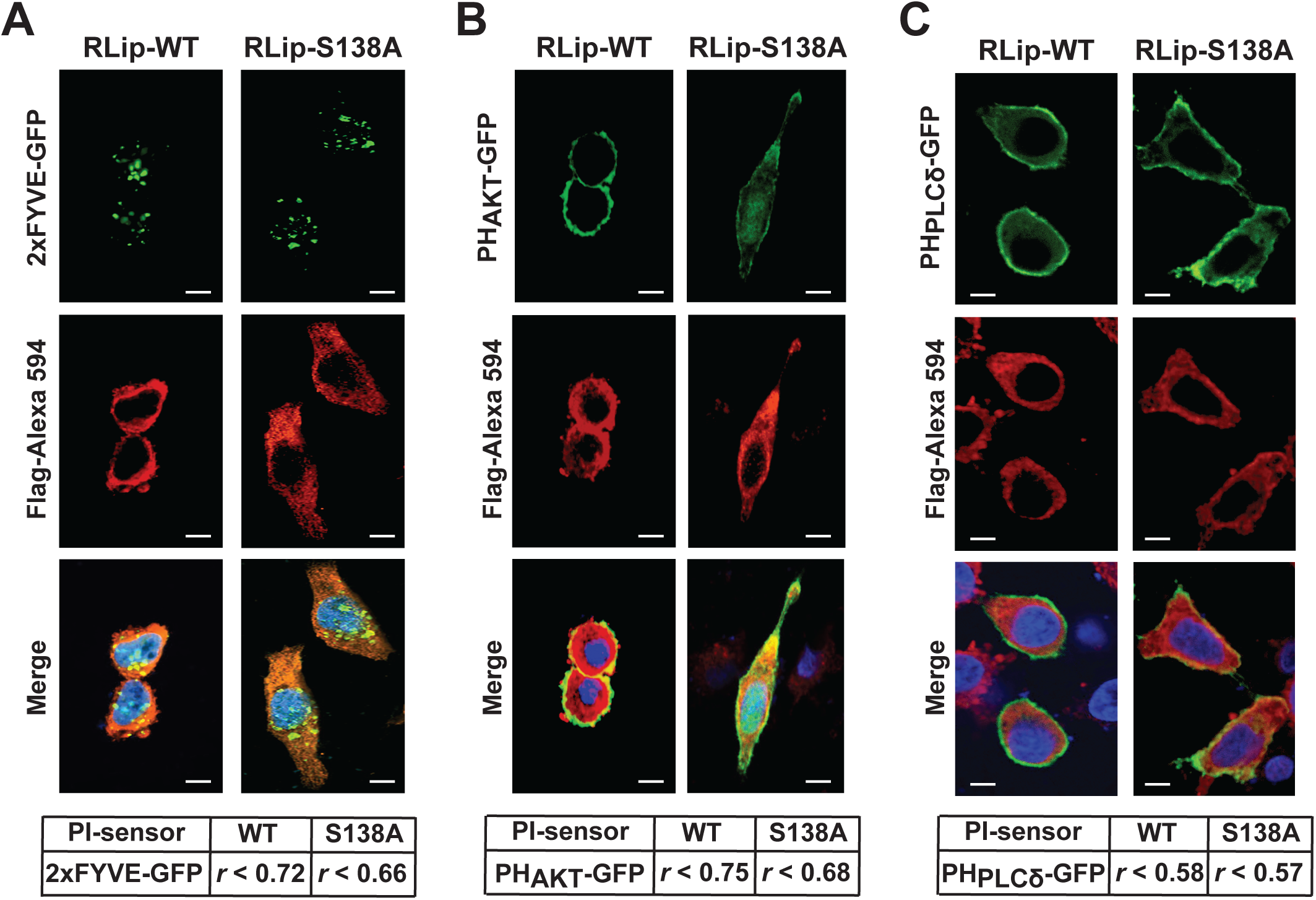
PI(3)P, PI(3,4,5)P_3_, and PI(3,4)P_2_ lipids are putative substrates of RLip. The pcDNA4-Flag-RLip-WT, or pcDNA4-Flag-RLip-S138A mutant constructs were co-transfected into HeLa cells with fluorescence PI probes for PI(3)P (GFP-2xFYVE) (**A**), PI(3,4,5)P_3_ and PI(3,4)P_2_ (GFP-PH_AKT_) (**B**), and PI(4,5)P_2_ (GFP-PH_PLCδ_) (**C**). Cells were fixed with 4 % PFA, and DNA was stained using 4’,6-diamidino-2-phenylindole (DAPI). Degree of colocalization between of RLip-WT or RLip-S138A (visualized via Alexa 594-conjugated Flag Ab staining) with the PI probes was analyzed using the Coloc 2 plugin from Fiji software and displayed as Pearson correlation coefficient (*r*)(58); 0 < *r* < 0.19 very low correlation 0.2 < *r* < 0.39 low correlation; 0.4 < *r* < 0.59 moderate correlation; 0.6 < *r* < 0.79 high correlation; 0.8 < *r* < 1 very high correlation. Bars, 10 μm. Images are a representative of 3 independent experiments.

### RLip facilitates intracellular survival of rickettsiae by promoting the evasion from lysosomal distraction

To test the biological role of RLip during rickettsial host invasion, we perform an antibody-mediated RLip neutralization assay. We first assessed the percentage of host cells infected by partially purified *R. rickettsii* pre-treated with anti-RLip antibody(ab) with respect to cells treated with pre-immune IgG. Our immunofluorescent antibody assay (IFA) showed a moderate, but significant, decrease in host internalization with cells pre-treated with anti-RLip Ab as compared to that of pre-immune IgG treated cells (**Figs. 4A**, and **4B**). These intriguing findings suggest that, in contrast to the well-described rickettsial Pat1 and Pat2 functional roles(19, 20, 23), RLip likely aids at later stages of the rickettsiae invasion process. Next, we evaluated the role of RLip during the phagosomal escape of rickettsiae into the host cell cytoplasm. In this effort, we also infected host cells with partially purified *R. rickettsii* pre-treated with either anti-RLip Ab or pre-immune IgG and assessed rickettsial phagosomal escape by IFA using the lysosomal marker, LAMP2 (lysosomal-associated membrane protein 2). We observed that *R. rickettsii* pretreated with anti-RLip Ab mostly colocalized with the LAMP2 marker, suggesting the rickettsiae remain enclosed within phagosomes (**Figs. 4C**, and **4D**). In contrast, *R. rickettsii* treated with pre-immune IgG did not colocalize with LAMP2 suggesting successful escape from lysosomal destruction (**Figs. 4C**, and **4D**). In sum, our data suggest that antibody-mediated neutralization of RLip function abrogates the successful escape of *R. rickettsii* from phagolysosomal destruction into host cytoplasm for colonization.

**Fig. 4.**
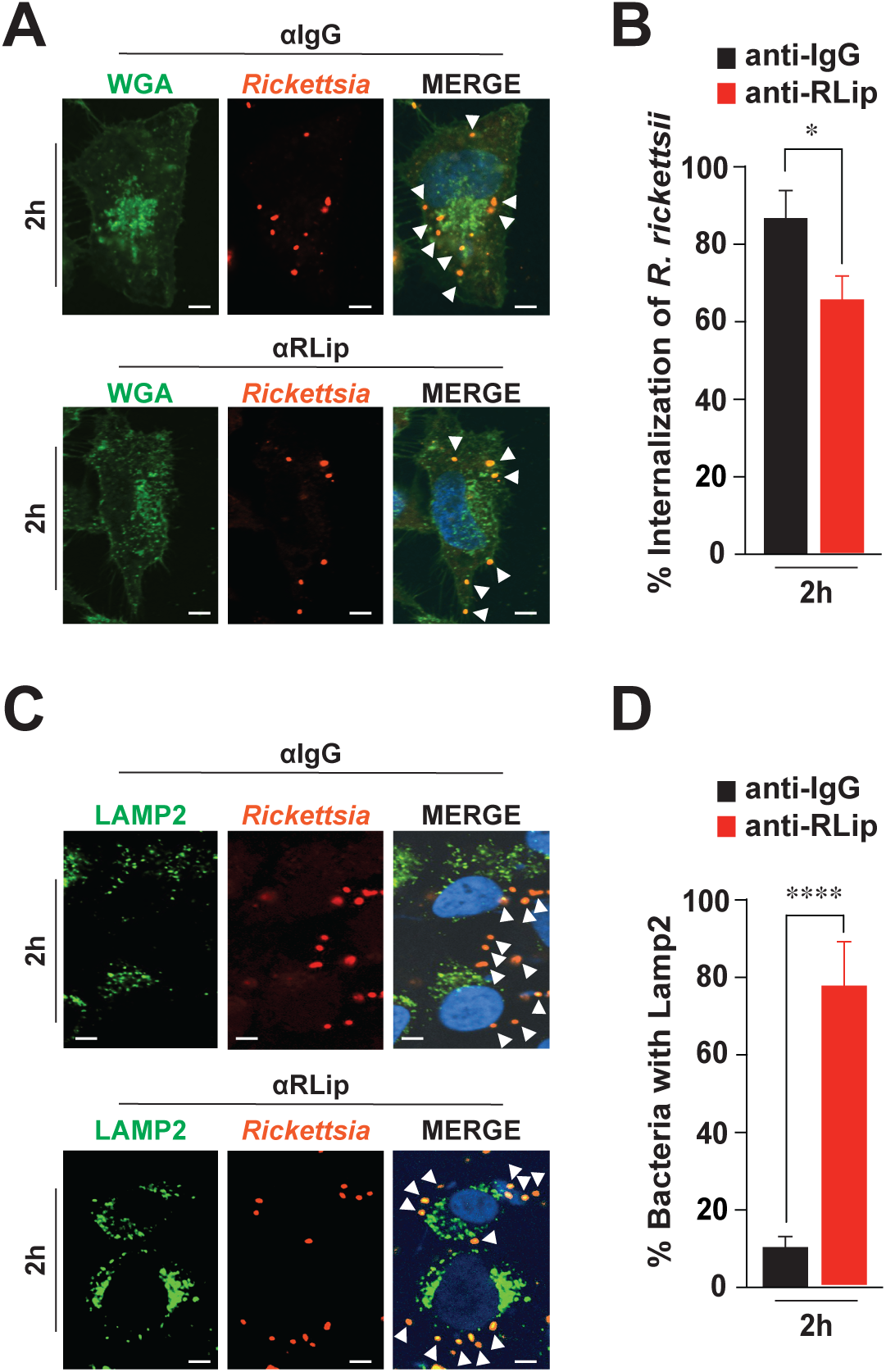
Antibody-mediated neutralization of RLip effects phagosomal escape of *R. rickettsii* (**A-B**) Partially purified *R. rickettsii* were pre-treated with 50 µg of affinity purified anti-RLip, or pre-immune IgG for 30 min on ice. Pretreated rickettsiae were added onto HMEC-1 monolayer followed by an additional 2 hrs incubation period at 34°C and 5 % CO_2_. Cells were fixed with 4 % PFA and incubated with Alexa Fluor-488-conjugated wheat germ agglutinin (WGA) (**A**), anti-*Rickettsia* guinea pig serum (**A**, and **C**), or anti-LAMP2 (**C**) Abs. The anti-guinea pig-Alexa Fluor-594 or anti-Alexa Fluor-488 were used as secondary Abs. The cell nuclei were stained with 4’,6-diamidino-2-phenylindole (DAPI). Colocalization between *Rickettsia* (highlighted by arrowheads) and WGA (**A**) or Lamp2 (**C**) was analyzed using Coloc 2 plugin Fiji software. Bars in panels **A**, and **C**, 10 μm. Images show *Rickettsia*-infected HMEC-1 cells after 2 hrs post-infection and are a representative of 3 independent experiments. Graphs show the percentage of *Rickettsia* internalization (**B**) or Lamp2 positive stained *Rickettsia* (**D**) at 2 hrs post-infection. Approximately 100 - 200 bacteria were counted per condition and time point. Error bars (**C**, **D**) represent means ± standard error of the mean (SEM) from three independent experiments; \**P* ≤ 0.05; \*\*\*\**P* ≤ 0.001.

## Discussion

*Rickettsia* species, overcome host defense responses to establish a successful intracytosolic replication niche. In fact, evasion of phagolysosomal destruction is essential for rickettsial intracellular survival and is mediated by their membranolytic effectors(19, 20, 22, 23, 25, 26). However, some of the rickettsial membranolytic enzymes have been reported to be dispensable depending on the host cell type, while others are sporadically present in rickettsial genome(21–23, 37), suggesting the presence of alternative membranolytic enzymes required for disrupting the vacuolar membrane to mediate escape from lysosomal destruction during host invasion. In this study, our bioinformatical analysis on the rickettsial genomes using the web-based Phyre2 program(24), identified a putative secretory lipase with a Serine hydrolase motif (GXSXG), named RLip. The secretory RLip molecule is highly conserved among all *Rickettsia* genomes, suggesting the potential relevance of RLip in rickettsial biology. The presented data here, showed RLip expression increased as early as 15 min post-infection and remained elevated until 2 hrs, indicating that RLip acts during an early stage of invasion.

The delivery of bacterial effectors involves various types of secretion pathways(37). For instance, the highly cytotoxic ExoU lipase (*P. aeruginosa*) is secreted via a type III secretion system (T3SS)(12, 38), while the VipD and VpdC lipases, from *Legionella pneumophila*, are secreted via the I-like (Dot/Icm) type IV secretion system (T4SS) (14, 15, 39). The secretion of rickettsial effectors involve various types of secretion pathways, which include the Sec translocon dependent type V secretion system (T5SS), type I secretion system (T1SS), type IV secretion system (T4SS) and others(21, 37, 40, 41). Intriguingly, analysis of RLip protein sequence by the web-based SignalP-6.0 program(42) did not predicted a N-term. signal sequence required for protein secretion via Sec translocon. Furthermore, the RLip protein was not captured by a RvhD4 coimmunoprecipitation assay performed in our prior reporting(32), suggesting that the molecule is not an effector of type IV secretion system (T4SS). However, using cellular fractionation assays, we demonstrated that RLip was secreted into the host cytoplasm during infection and was not retained by the bacteria itself. These findings would suggest that either the T1SS or any other unappreciated protein secretion pathways could be involved in the secretion process of RLip. To our knowledge these data also provide the first description of a rickettsial effector that is synthesized during infection, secreted into host cytoplasm and retained minimally by the bacteria itself during host cell-free stage. Intriguingly, RLip expression and secretion patterns shows similarities to the chlamydial protease- or proteasome-like activity factor (CPAF)(43) and hypothetical protein CT795(44). In fact, CPAF, like RLip, is secreted during the infection process and remains hardly associated with purified *Chlamydia* and therefore was designated as an infection-dependent antigen(43, 44). Thus, it is tempting to speculate that RLip may acts as a secreted rickettsial infection-dependent antigen, which will be addressed in our future work.

Bacterial membranolytic effectors have been shown to play a critical role in mediating membrane ruptures, during infection, that includes *Pseudomonas aeruginosa* (ExoU)(12, 13), *Listeria monocytogenes* (PLCs)(7–11), *Legionella pneumophila* (VipD and VpdC)(14, 15) and *Shigella flexneri* (IpaB and IpaC)(16–18). In addition, viruses use lipases to overcome endosomal membrane barriers such as parvovirus capsid protein VP1(45), or the picornaviruses recruited host lipid-modifying enzyme (PLA2G16)(46). In case of *Rickettsia*, work from other laboratories and ours have demonstrated that PLD(22, 25, 26), Pat1, and Pat2(19, 20, 23) are important membranolytic enzymes. Here, we are reporting that *R. rickettsii* RLip effector has a catalytic active site Serine at position 138, required for its lipase enzyme activity and cytotoxicity. Moreover, we found that the RLip lipase activity was increased in the presence of host cell lysates suggesting the involvement of a host factor(s) to promote its biological activity during host invasion. These findings agree with earlier reports that showed eukaryotic factor(s) enhanced the enzymatic activities of both rickettsial lipases, Pat1 and Pat2(19, 20, 23). However, the identity of those eukaryotic co-factor(s) remains unknown, which warrants further investigation. The lower lipase enzymatic activity of RLip, that is further strengthened by eukaryotic factor(s), may be necessary to support the obligate intracellular lifestyle of rickettsiae, without inflicting any rapid damage to host cells, at least before rickettsiae exit infected cells to promote further transmission to neighboring cells/organs of the host.

Rickettsial membranolytic effectors play a critical role in creating a habitable intracellular niche, however, their biological relevance in aiding the infection process depend on the host cell-type and the specific *Rickettsia* species(21–23, 37). For instance, Pat2 is sporadically present in rickettsial genomes, while Pat1 is highly conserved in all *Rickettsia* species(21, 37). Intriguingly, Pat1 was shown to be redundant for rickettsial growth in HMEC-1, however, required for survival in bone-marrow derived macrophages (BMDM)(23). To test our hypothesis that *Rickettsia* requires additional enzyme(s) to escape from vacuolar membrane, our bioinformatic search identified a new membranolytic enzymes, RLip, which is highly conserved in all *Rickettsia* species. We further assessed RLip’s role during live infection by employing an antibody-mediated neutralization assays and demonstrated that pre-treatment of *R. rickettsii* with anti-RLip Ab only modestly affected rickettsiae host internalization, however, it strongly impaired the escape from lysosomal fusion. Intriguingly, preceding findings from our laboratory showed that pre-treatment of *R. typhi* with either anti-Pat1 or anti-Pat2 Ab blocked rickettsial infection as well as delayed rickettsial phagosome escape(20), suggesting that RLip has a distinct functional role during host invasion. Our cellular localization assays using selective PI biosensors revealed a strong colocalizations of RLip with phosphoinositides that are mostly found during the early events of the endocytic pathways [PI(3,4)P_2_], including early endosomes [PI(3)P](32, 35, 36). Taken together, these data suggests that membranolytic effector RLip plays a role in facilitating the escape into host cytoplasm by evading the trafficking to the lysosomes.

It is important to note that rickettsial vacuolar escape from lysosomal fusion into host cytosol involves is a multi-effector process. For instance, another potential protein involved in rickettsial escape from phagolysosomal fusion could be hemolysin (TlyC)(25, 47), which exhibit similar functions as listeriolysin O (LLO) of *Listeria monocytogenes*, a cholesterol-dependent cytolysin(7, 8, 48). Recently, we identified a rickettsial PI3-Kinase, Risk1, that interacted with PI(3)P to facilitate bacterial escape by delaying the vacuolar fusion with destructive lysosome(32). Collectively, based on the data presented in this manuscript it is tempting to speculate that RLip alone or in combination with other rickettsial effector(s) facilitates successful escape from vacuolar membranes.

## Materials and Methods

### Antibodies and reagents

Anti-LAMP2 (H4B4), anti-GAPDH (FL-335), and horseradish peroxidase (HRP)-conjugated secondary Abs (mouse, rabbit, rat, guinea pig, and goat IgGs) were purchased from Santa Cruz Biotechnology. ProLong Gold antifade mounting medium with DAPI (4′,6-diamidino-2-phenylindole), paraformaldehyde (PFA), Halt protease and phosphatase inhibitor cocktail, HisPur^TM^ Ni-NTA magnetic beads, anti-V5 (SV5-Pk1), anti-His (C-term., 46-0693) Ab, and Alexa 488/594-conjugated secondary Abs and wheat germ agglutinin (WGA) were purchased from Thermo Fisher Scientific. Anti-Flag (M2) Ab was purchased from Sigma, while the PolyJet transfection reagent was obtained from Signagen.

### Bacterial strains, cell culture, and infection

Vero76 (African green monkey kidney, RL-1587; ATCC), SVEC4-10 (CRL-2181, ATCC), and HeLa (CCL-2; ATCC) cells were maintained in minimal Dulbecco’s modified Eagle’s medium (DMEM) supplemented with 10% heat-inactivated fetal bovine serum (FBS) at 37°C with 5% CO_2_. HMEC-1 cells (human microvascular endothelial cells, CDC, Lot-No.: 119223) were grown in HMEC-1 media [MCDB 131 media (Invitrogen) supplemented with 10 % FBS (GeminiBio), 10 mM L-glutamine (Gibco), 10 ng/ml epidermal growth factor (Becton-Dickinson), 1 μg/mL hydrocortisone (Sigma), and 1.18 mg/mL sodium bicarbonate]. *R. rickettsii* (Shelia Smith) strain was obtained from Dr. Ted Hackstadt (Rocky Mountain Laboratories, NIH, MT, USA). *R. parkeri* (Portsmouth) and *R. montanensis* strains were obtained from the CDC. All *Rickettsia* species were propagated in Vero76 cells grown in DMEM supplemented with 5 % FBS at 34°C with 5 % CO_2_. *R. rickettsii*, *R. parkeri*, and *R. montanensis* were partially purified as described previously(49, 50). Briefly. rickettsiae infected cells were disrupted by mild sonication for 5 sec using a sonic dismembrator (Fisher Scientific). The disrupted host cells were centrifuged at 300 x *g* for 5 min at 4°C to remove host cell debris or any remaining intact host cells. The supernatant was filtered through a 5-μm pore size filtering unit (Millipore). The filtrate was further centrifuged at 10,000 x *g* for 5 min at 4°C. The pellet containing partially purified rickettsiae were collected for host cell infection and RLip experiments. For early stages of infection [before the doubling time (8 to 10 hrs) of rickettsiae], a higher multiplicity of infection (MOI) of 20 [for 0.5 and 2 hrs post-infection (hpi)] was used to ensure the presence of sufficient number of bacteria, as compared to MOI of 5 at later time points (24 hpi), to determine the biological functions of the bacteria during host infection(19, 32, 50, 51).

### Bioinformatic analysis and Homology modelling of RLip

The sequence analyses of RLip using blastp (against the NCBI Conserved Domains Database)(52) and Phyre2(24) suggests that RLip possesses a secretory lipase with Serine hydrolase motif (GXSXG). The putative active site region of RLip (locus_tag: A1G_01170) from *R. rickettsii* (Sheila Smith) strain was aligned with other bacterial lipases [*Pseudomonas aeruginosa* ExoU (WP_003134060); *Legionella pneumophila* VipD (WP_010948518); *Legionella pneumophila* VpdC (WP_010947155); *R. typhi* Pat1 (WP_011191036), and *R. typhi* Pat2 (AAU03991)]. The sequence alignment of RLip homologs across other rickettsiae [*R. bellii* (WP_012151906); *R. akari* (ABV74559); *R. australis* (WP_014413153); *R. felis* (WP_041405363); *R. montanensis* (WP_014409937); *R. conorii* (WP_010976872); *R. rickettsii* (WP_012150422); *R. parkeri* (WP_014410398); *R. canadensis* (WP_012148317); *R. typhi* (WP_011190628); and *R. prowazekii* (WP_004598629)] was done by Clustal Omega using default parameters. The homology modelling of RLip was performed by automated sever Phyre2(24). The model was validated by Ramachandran plot using RAMPAGE(31) and visualized by PyMOL (Molecular Graphics System, Schrödinger, LLC). Further analysis of the RLip protein sequence using the web-based SignalP - 6.0 program(42) showed no presence of a signal peptide sequence required for Sec translocon.

### Antibody against rickettsial antigen

The rabbit antibody against recombinant full-length RLip protein (anti-RLip), encoded by codon-optimized A1G_01170 (locus_tag) of *R. rickettsii* (Sheila Smith) strain, was generated and affinity purified by Thermo Fisher Scientific. The rabbit antibody against recombinant Pat1 protein [283 amino acids of the C-term. region), encoded by locus_tag: A1G_05085] (anti-Pat1) of *R. rickettsii* (Sheila Smith) strain, was generated and affinity purified by Thermo Fisher Scientific. The mouse monoclonal antibody against SFG rickettsiae, anti-OmpA/B (clone: RC-5H2) was purchased from Fuller laboratories. The specificity of all antibodies was validated by immunoblotting (please see **Fig. S2B**, and **S2C**).

### Mammalian expression plasmids

The codon-optimized full-length RLip-WT and RLip-S138A mutant were sub-cloned into green fluorescent protein tagged (pcDNA6.2), and FLAG-tagged (pcDNA4/TO/StrepII) plasmid. All the constructs were confirmed by sequencing. pGFP-2xFYVE was a kind gift from George Banting(53). The pGFP-PH_AKT_ plasmid was a kind gift from Craig Montell (Addgene plasmid 18836)(54). The pGFP-PH_PLCδ_ construct (Addgene plasmid 21179) was kindly gifted by Tobias Meyer(55).

### Secretion assay

Monolayer of Vero76, or HMEC-1 cells, either infected or uninfected with *R. rickettsii*, were incubated in culture medium at 34°C as described elsewhere(20, 32). Briefly, cells were lysed in 1 x PBS buffer (containing 0.1% Triton X-100, protease and phosphatase inhibitors) for 15 min on ice(56). Lysates were centrifuged for 5 min at 6,000 x *g* to separate the rickettsial secreted effectors and host cytosolic proteins (cytoplasmic fraction) from the intact rickettsiae and insoluble host proteins (pellet fraction). The cytoplasmic fraction was filtered through a 0.45-μm pore size filter (Millipore), and pellet fraction was resuspended into PBS (containing protease and phosphatase inhibitors). Samples from the pellet (P) and cytoplasmic (C) fractions were immunoblotted with anti-RLip, anti-Pat1 (positive control for secreted *R. rickettsii* effector), anti-OmpA/B (as control for *R. rickettsii* surface protein), or anti-GAPDH Abs (host cytoplasmic control protein)(20, 32).

### Expression and purification of recombinant RLip proteins for lipase activity assay

The *R. rickettsii RLip* (*<u>R</u>. rickettsii* <u>Lip</u>ase, gene locus_tag # A1G_01170) gene was codon-optimized (CO) and cloned into the bacterial expression vector pET30a with N-term. 6x-His epitope tag by GeneScript. Mutation of the catalytic active site at position 138 (Ser-138 to Ala-138) of RLip was introduced via the QuikChange II XL site-directed mutagenesis kit (Agilent technologies, Cat-No.: 200521-5) according to the manufacturer’s instructions using the using Forward primer 5’-CTTTTAACACCAGCGGCCACGCGCTGGGCGCGATTATC-3’ and Reverse primer 5’-GATAATCGCGCCCAGCGCGTGGCCGCTGGTGTTAAAAG-3’. The constructed mutant plasmid pET30a-RLip-S138A (CO) was confirmed by sequencing. Both pET30a-RLip-WT (CO) and RLip-S138A (CO) plasmids were transformed into *E. coli* BL21-Codon Plus expression host and grown at 37°C for overnight in LB media supplemented with Kanamycin (30 µg/ml). Protein expression was induced using 0.5 mM IPTG for 16 hrs at 20°C. Proteins were purified using HisPur^TM^ Ni-NTA magnetic beads according to manufacturer’s instructions and identity of the recombinant proteins was confirmed by western blot analysis using an anti-His Ab (**Fig. 2A**) as well as by MS analysis (CVID core facility, University of Maryland School of Medicine, Baltimore, MD, USA).

### Lipase activity assay

The lipase activity of recombinant codon-optimized RLip-WT-CO and RLip-S138A-CO proteins was calorimetrically measured using a lipase assay kit (Abcam, Cat-No.: ab102524) following manufacturer’s instruction. Briefly, monolayers of HMEC-1 or Vero76 cells were harvested and washed by PBS. Cell pellets were resuspended in 0.5 ml of assay buffer, lysed by sonication and supernatant was collected by centrifugation at 10,000 x *g* for 15 min at 4°C. The protein concentration of supernatants was determined by BCA protein assay kit. Freshly purified recombinant (r)RLip-WT-CO and rRLip-S138A-CO proteins were buffer exchanged with the provided assay buffer. For lipase assay, approximately, 25 µg of rRLip-WT-CO and rRLip-S138A-CO proteins and 50 µg of either Vero76 or HMEC-1 cell lysates were mixed with reaction buffer containing lipase substrate, OxiRed probe, and enzyme mixture. Samples were mixed and measured at an OD_570 nm_ for 90 min, at intervals of 2 min, at 37°C. The Lipase activity of the rRLip-WT-CO and rRLip-S138A-CO proteins was calculated following manufacturer’s instruction.

### Yeast cytotoxicity assay

The *R. rickettsii* genome encoded wild-type *RLip* gene was cloned by PCR into the *SacI* and *XhoI* sites of yeast expression vector pYES2/CT with C-term. epitope (V5 and 6x-His) tags according to manufacturer’s instruction, using Forward primer 5’-TTAAGCTTGGTACCGAGCTCATGCCTACGTACAAAAATTCTAAACATATTAGCAC-3’ and Reverse primer 5’-GCCCTCTAGACTCGAGTTAACATATTAGAGGATATAGATGATAATTACTTATATCTCCTGTTGT-3’. Constructed plasmid pYES2/CT-RLip-WT_GS_ (carrying RLip encoded by *R. rickettsii* wild-type genome sequence (GS)] was confirmed by sequencing. Next, the codon-optimized (CO) *R. rickettsii RLip* gene (GeneScript) was subcloned by PCR into the *SacI* and *XhoI* sites of yeast expression vector pYES2/CT using Forward primer 5’-TTAAGCTTGGTACCGAGCTCATGCCCACATATAAAAATTCAAAGCACATATCC-3’ and Reverse primer 5’-GCCCTCTAGACTCGAGGTGGTGATGATGGTGGTGACAAATCAGCGG-3’ and constructed plasmid pYES2/CT-RLip-WT-CO was confirmed by sequencing.

In addition, codon-optimized (CO) catalytic active site mutant (S138A-CO) of RLip was subcloned into the *SacI* and *XhoI* sites of yeast expression vector pYES2/CT and constructed plasmid pYES2/CT-RLip-S138A-CO was confirmed by sequencing as described above.

The constructed plasmids were transformed in *S. cerevisiae* strain INVSc-1 using Frozen-EZ Yeast Transformation II^TM^ Kit (Zymo Research, Cat-No.: T2001) following the manufacturer’s instructions. The transformed yeast cells were grown in synthetic complete (SC) medium agar without Uracil and containing 2% glucose (SC-U+Glu) to select plasmids at 30°C for 3 days. Plasmid containing yeast cells were grown in SC-U+Glu media overnight at 30°C. The culture was pelleted, washed, and resuspended in SC-U without a carbon source. The resuspended yeast cells were induced and/or repressed in agar medium containing 2 % galactose (SC-U+Gal) and/or 2 % glucose (SC-U+Glu) respectively and incubated for 3 days at 30°C. For CFU assay, the resuspended yeast transformants were serially diluted in SC-U without a carbon source and plated on inducing (SC-U+Gal) and repressing (SC-U+Glu) agar. After incubation at 30°C for 3 days, the colonies were counted to determine the percentage of CFU on inducing agar with respect to that on repressing agar.

### Immunofluorescent assay (IFA)

Eight-well chamber slides were seeded with HeLa cells (∼ 50 x 10^4^ cells per well) and infected with partially purified *R. rickettsii* (MOI = 20 [2 hrs] as described previously(32, 50, 51, 57). Briefly, rickettsiae were added to HeLa cells and incubated for 2 hrs at 34°C with 5 % CO_2_. Following incubation, cells were washed with 1 x phosphate-buffered saline (PBS) and fixed with 4 % paraformaldehyde (PFA) for 20 min at room temperature. Cells were then permeabilized in blocking buffer (0.3 % saponin and 0.5 % normal goat serum in 1 x PBS) for 30 min and incubated for 1 hr with the following primary antibodies diluted in Ab-dilution buffer (0.3 % saponin in 1 x PBS): anti-*Rickettsia* (1:100, guinea pig serum), and anti-Lamp2 (1:100). Cells were then washed with 1 x PBS and incubated for 1 hr with anti-Alexa Fluor 488 or anti-Alexa Fluor 594 secondary Abs diluted 1:1,500 in Ab-dilution buffer or Alexa Fluor 488-conjugated wheat germ agglutinin (WGA). Next, cells were washed with 1 x PBS and mounted with ProLong Gold antifade mounting medium containing DAPI. Images were acquired using the Nikon W-1 spinning disk confocal microscope (University of Maryland School of Medicine, Confocal Core Facility, Baltimore, MD, USA) and colocalization strength between *Rickettsia*, and WGA or Lamp2 was analyzed using Fiji software as described previously(32, 51, 57). The percentage of internalized bacteria (approximately 200 bacteria were counted per strain and time point) was calculated by dividing the number of extracellular bacteria by the total number of bacteria, multiplying by 100, and then subtracting this number from 100 % to get the percentage of intracellular bacteria.

For experiments investigating the effects of RLip on the distribution of various cellular phosphoinositides, HeLa cells were transfected with 0.5 μg of pcDNA4/TO/StrepII-Flag empty vector, pcDNA4/TO/StrepII-Flag-RLip-WT, or pcDNA4/TO/StrepII-Flag-RLip-S138A plasmid DNA, in combination with 0.5 μg of pGFP-PH_PLCδ_, pGFP-2xFYVE, or pGFP-PH_AKT_ plasmid DNA using PolyJet transfection reagent (Signagen) per the manufacturer’s recommendation. Changes in phosphoinositide distribution were monitored in cells that co-expressed RLip constructs (visualized via Alexa 594-conjugated Flag Ab staining) with the corresponding phosphoinositide probes. Approximately 200 co-transfected cells were enumerated by confocal microcopy, and each experiment was repeated in triplicates. Colocalization strength between phosphoinositide probes and RLip-WT or RLip-S138A was analyzed using the Coloc 2 plugin for Fiji software and displayed as Pearson correlation coefficient (*r*)(58); 0< *r* <0.19 very low correlation 0.2< *r* <0.39 low correlation; 0.4< *r* <0.59 moderate correlation; 0.6< *r* <0.79 high correlation; 0.8< *r* <1 very high correlation.

### Antibody-mediated neutralization of RLip

Partially purified rickettsiae were treated with 50 µg of affinity purified anti-RLip, or pre-immune IgG for 30 min on ice. Pretreated rickettsiae were added onto Hela cells followed by centrifugation at 200 x *g* for 5 min at room temperature to induce contact and incubated for 2 hrs at 34°C. Cells were washed with 1 x PBS and fixed with 4 % PFA at room temperature for 20 min. The cells were immunolabeled and analyzed as described under IFA.

### RNA isolation and quantitative real-time PCR

Rickettsial growth [assessed by rickettsial citrate synthase (*GltA*) gene expression] and RLip gene expression was determined after *R. rickettsii* infection in various cell lines (Vero76, HMEC-1, and SVEC4-10) by real-time reverse transcription-quantitative PCR (RT-qPCR)(19, 51, 57). In this effort, host cells were infected with *R. rickettsii* and collected at 0.25, 0.5, 1, 2, 24, 48 and 72 hrs post-infection. RNA was isolated from ∼ 1 x 10^6^ cells using Quick-RNA^TM^ miniprep kit (Zymo research, Cat-No.: ZR1055). cDNA was synthesized from the RNA samples using GoScript™ Reverse Transcription System (Promega) as per manufacturer’s instructions. After cDNA synthesis, amplification reaction was carried out by iQ^TM^ SYBR^®^ Green Supermix (Bio-Rad), using Forward primer 5’-GACTTGCTTATCGCACTGCAC-3’ and Reverse primer 5’-TGCTCCAAGTGAATGACCGCT-3’ for RLip. Expression of host cell housekeeping gene, *GAPDH*, and rickettsial *GltA*, was determined as described previously(51, 57). Cycling conditions were as follows: 1 cycle at 95°C for 3 min; 40 cycles at 95°C for 15 sec, 55°C for 15 sec, and 72°C for 20 sec; and 1 cycle to generate the dissociation curve. The RT-qPCR amplification and detection were performed on an QuantStudio™ 3 Real-Time PCR Systems (Applied Biosystem by Thermo Fischer). The specificities of these primer pairs were verified by PCR on DNA isolated from *R. rickettsii*. Rickettsial growth (*GltA*) and *RLip* expression were normalized with respect to *GAPDH* expression. The relative copy number was calculated with the equation *RCN= E^-Δct^*, where *E* = efficiency of PCR and ΔCt = Ct *target* − Ct *GAPDH* as described previously(51, 57).

### Extract preparation and Western blot analysis

*Rickettsia*-infected Vero76, HMEC-1, SVEC4-10, or HeLa cells were lysed for 2 hrs at 4°C in ice-cold lysis buffer (50 mM HEPES [pH 7.4], 137 mM NaCl, 10% glycerol, 1 mM EDTA, 0.5% NP-40, and supplemented with protease and phosphatase inhibitory cocktails) as described previously(32). Equal amounts of protein were loaded onto an SDS-PAGE and membranes were probed with anti-RLip, anti-Pat1, anti-V5, anti-His, anti-GAPDH, and anti-OmpA/B Abs, followed by enhanced chemiluminescence with secondary Abs conjugated to horseradish peroxidase.

### RLip induction assay

HMEC-1 cells were infected for 0.5 and 2 hrs with partially purified *R. rickettsii* from host cells. Samples were lysed in ice-cold lysis buffer supplemented with protease and phosphatase inhibitory cocktails and equal amounts of protein were loaded onto an SDS-PAGE. Membranes were probed with anti-RLip, anti-GAPDH, and anti-OmpA/B Abs, followed by enhanced chemiluminescence with secondary Abs conjugated to horseradish peroxidase.

### Statistical analysis

The statistical significance was assessed using analysis of variance with Bonferroni’s procedure and Student’s *t*-test. Data are presented as mean ± standard error of the mean (SEM), unless stated otherwise and asterisks denote statistical significance as: \**P* ≤ 0.05; \*\**P* ≤ 0.01; \*\*\**P* ≤ 0.001; \*\*\*\**P* ≤ 0.0001, compared with the indicated controls. Statistical analyses were performed using GraphPad PRISM, version 9.

## Acknowledgments

We thank George Banting for pGFP-2xFYVE plasmid and Tobias Meyer (Stanford University, CA, USA) for the pGFP-PH_PLCδ_ (Addgene 21179) plasmid, Craig Montell for the pcDNA3-GFP-PH_AKT_ plasmid (Addgene plasmid 8836). We are also grateful to Ted Hackstadt (Rocky Mountain Laboratories, NIH, MT, USA) for generously providing us with essential biological reagents, including antibodies and *Rickettsia* strains. Sincerely, we acknowledge Dr. Edwin Ades and Mr. Fransisco J Candal of CDC and Dr. Thomas Lawley of Emory University as the developers of HMEC-1 (human microvascular endothelial cells).

We also would like to thank Abdu F. Azad and Magda Sexton (University of Maryland School of Medicine, Baltimore, MD, USA) for their support and guidance during design and planning of this RLip project.

This work was supported with funds from the NIAID/NIH grants (R01AI017828 and R01AI126853 to M.S.R., and R21AI166821 to O.H.V. and M.S.R.).

The funding sources had no role in the design of the study, in the collection, analyses, or interpretation of data, in the writing of the manuscript, or in the decision to publish the results.

## Supplementary figure legends

**Fig.S1. RLip harbors a classical lipase structure.**

**(A)** Homology model of RLip, showing a classical lipase structure of a β-sheet surrounded by two α-helices, was constructed using the Phyre2 software(30). (**B**) Homology models of lipase structures from other bacterial phospholipases (ExoU, VipD, VpdC, and rickettsial Pat1, or Pat2) was constructed as described above.

**Fig.S2. RLip is rapidly induced and secreted during *R. rickettsii* infection of Vero76 cells.**

**(A)** Expression kinetics of *RLip* (black line) and rickettsial citrate synthase (*GltA,* red line) during *R. rickettsii-* infection of Vero76 cells was measured by RT-qPCR. Rickettsial growth was determined by *GltA* expression. Both *RLip* and *GltA* expression were normalized with respect to *GAPDH* expression as described previously(51, 57). (**B**) Codon-optimized RLip protein was used to generate an anti-RLip Ab (RLip). The specificity of the anti-RLip Ab was validated by western blot analysis using lysates of uninfected and *R. rickettsii*-infected Vero 76 cells [lanes 1 (Uninfected) and 2 (72 hrs post-infection)] and recombinant (r)RLip-WT protein (lanes 3 to 5; 10-500 ng). Immunoblotting with anti-OmpA/B and anti-GAPDH Abs was used as controls of the uninfected and *R. rickettsii*-infected Vero 76 cells (lanes 1 and 2). (**C**) Uninfected or *R. rickettsii*-infected Vero76 cells were lysed with 0.1 % Triton X-100 and separated into cytoplasmic (C) and pellet (P) fractions. Samples were immunoblotted with anti-RLip, anti-Pat1 (positive secreted *R. rickettsii* effector control), anti-OmpA/B (as control for *R. rickettsii* surface protein), or anti-GAPDH Abs (host cytoplasmic control protein). Whole cell lysates (WCL) were used as expression control for all target proteins. Error bars in panel **A** represent means ± standard error of the mean (SEM) from three independent experiments. Images in panels **B**, and **C** are a representative of 3 independent experiments.

**Fig. S3. Immunofluorescent staining of HeLa cells co-transfected with empty vector and PI probe plasmids.**

pcDNA4-Flag empty vector plasmid was co-transfected into HeLa cells with fluorescence PI probes for PI(3)P (GFP-2xFYVE) (**A**), PI(3,4,5)P_3_ and PI(3,4)P_2_ (GFP-PH_AKT_) (**B**), and PI(4,5)P_2_ (GFP-PH_PLCδ_) (**C**). Cells were fixed with 4 % PFA, and DNA was stained using 4’,6-diamidino-2-phenylindole (DAPI). Degree of colocalization between of empty vector (using Alexa 594-conjugated Flag Ab staining) with the corresponding PI probes. Approximately 200 co-transfected cells were enumerated by confocal microcopy, and each experiment was repeated in triplicates. Colocalization between PI probes and RLip-WT or RLip-S138A was analyzed using the Coloc 2 plugin for Fiji software and displayed as Pearson correlation coefficient with the PI probes was analyzed using the Coloc 2 plugin from Fiji software and displayed as Pearson correlation coefficient (*r*)(58); 0 < *r* < 0.19 very low correlation. Bars, 10 μm. Images are a representative of 3 independent experiments.

## Author Contributions

M.S., M.S.R., and O.H.V. planned the research, analyzed, and interpreted the data; M.S., I.M., S.U., O.H.V., and M.S.R. performed the experiments; M.S.R. and O.H.V. contributed to the overall project administration and supervision; M.S., M.S.R., and O.H.V. wrote the manuscript; and all authors participated in editing the manuscript.

